# Cerebrovascular Claudin-5 Isoform Expression Correlates with Worsened Stroke Outcomes Following Thromboembolic Stroke

**DOI:** 10.1101/2025.08.13.670217

**Authors:** Trevor S. Wendt, Henrik Andersson, Kajsa Arkelius, Saema Ansar

**Affiliations:** Applied Neurovascular Research, Neurosurgery, Department of Clinical Sciences, Lund University, Klinikgatan 28, BMC C12, 222 42, Lund, Sweden

**Author notes:** **Corresponding Author**: Saema Ansar.

## Abstract

**Background and Purpose:** Claudin-5 plays a crucial role in the maintenance of the blood-brain barrier (BBB) integrity through its role in endothelial tight junction formation. Alternative splicing of claudin-5 within the microvascular endothelium may modulate BBB structural and functional dynamics, potentially influencing neuronal damage and recovery following ischemic stroke. We hypothesized that ischemic stroke induces temporal changes in claudin-5 protein isoform expression that correlates with worsened neurological outcomes.

**Methods:** Male Wistar rats underwent thromboembolic stroke. Claudin-5 isoform expression was assessed at 3, 6, and 24h post-stroke onset, with additional groups receiving recombinant tissue plasminogen activator (rt-PA) at 4 hours post-stroke. Brain edema, infarct volume, hemorrhage, and cerebral blood flow was evaluated using 9.4T MRI. Ipsilateral and contralateral cerebrovascular claudin-5 expression was quantified via western blotting while neurological function was assessed by 28-point neuroscore. In addition, RNA sequencing analysis was performed to identify novel splice variants.

**Results:** A time-dependent increase in claudin-5 isoform 1 (35kDa) expression levels in the ipsilateral cerebrovasculature at 6 h was observed. Isoform 2 (25kDa) and fragment (10kDa) isoforms of claudin-5 remain unchanged. Treatment with rt-PA maintained the elevated levels of isoform 1 claudin-5 protein expression within the ipsilateral hemisphere. Increased claudin-5 isoform 1 expression within the ipsilateral hemisphere correlated with increased brain edema, hemorrhage, and worsened neurological function at 24h post-stroke onset. RNA sequencing revealed novel *CLDN5* splice isoforms in post-stroke rat brain tissue which resemble structural similarity to known human *CLDN5* isoforms.

**Conclusion:** These findings demonstrate that ischemic stroke induces temporal, hemisphere-specific alterations in claudin-5 isoform expression that correlate with BBB dysfunction and poor neurological outcomes. The potential indication of novel alternative splice variants suggests that post-transcriptional regulation of claudin-5 represents a previously unrecognized mechanism contributing to endothelial tight junction dysfunction and stroke pathophysiology. These results highlight claudin-5 isoform expression as a potential therapeutic target for preserving BBB integrity following cerebral ischemia.

## Introduction

The blood-brain-barrier (BBB) is a highly specialized, selective, and conserved interface that is critical to the maintenance of central nervous system (CNS) homeostasis by tightly regulating molecular exchange between the blood and parenchymal tissue.^1–5^ There are many components that contribute to the integrity and functionality of the BBB including astrocytes, pericytes, microglia, and the endothelium.^6–9^ While all these components play crucial roles in the physiologic maintenance of the BBB, the microvascular endothelium have emerged as the structural core upon which the rest of the components rely upon to carry out their functions.^10,11^ Within the endothelium lies a concentrated network of intercellular complexes including transmembrane and cytoplasmic proteins which forms a series of interconnected cells that establishes this foundation.^12^ This network is mediated primarily via tight junctions such as occludin, junctional adhesion molecules, and claudins.^13^ Among these, claudin-5 is the most abundantly expressed and functionally dominant tight junction protein within the cerebral microvasculature, which is essential for maintaining the low paracellular permeability that defines the neurovascular unit.^14,15^ Unlike other claudins, claudin-5 is uniquely enriched within the cerebrovascular endothelium, and its spatial distribution and tight junctional localization are tightly regulated to sustain the integrity of the barrier under physiological and pathophysiological conditions.^16,17^ While foundational work has demonstrated the necessity of claudin-5 in restricting small molecule passage through the BBB^18^, less is known about how claudin-5 expression is dynamically regulated in response to injury, particularly at the transcriptional and post-transcriptional levels.

This is exemplified by the current understanding that claudin-5 is traditionally expressed as a single isoform protein; however, emerging evidence suggests the existence of alternative splice variants^17^ which may play a role in progression of neurological disease states.^19^ A study identified two common human *CLDN5* alleles encoding different open reading frames, resulting in 218 and 303 amino acid proteins.^20,21^ However, to date only the 218 amino acid isoform has been detected in human tissues, indicating a potential predominant expression of this variant under physiological conditions. These isoforms may lead to unique differential localization within the endothelium, integration into tight junction strands, and responses to pathological stimuli such as acute ischemic stroke. However, a thorough investigation into claudin-5 isoform expression in the context of ischemic stroke remains to be elucidated, which might yield novel mechanistic insight into the pathophysiological progression of ischemic stroke.

Cerebrovascular injury such as ischemic stroke and traumatic brain injury is tightly associated with BBB breakdown and is a leading cause of morbidity and mortality worldwide.^22–26^ BBB disruption exacerbates edema, hemorrhagic transformation, and secondary neuronal injury, culminating in worsened neurological outcomes. Although numerous studies have focused on changes in BBB permeability following stroke, the molecular underpinnings mediating endothelial barrier compromise, particularly at the level of tight junction isoform regulation, remain incompletely defined. The potential emergence or temporal fluctuation of distinct claudin-5 isoforms in response to cerebral ischemia represents a critical knowledge gap that could inform novel therapeutic approaches for BBB preservation and neurological recovery. To address this limitation, we conducted a comprehensive investigation of the temporal expression of claudin-5 isoforms following experimental thromboembolic stroke and determined their relationship to BBB integrity and neurological outcomes. Specifically, we aimed to: (1) characterize alterations in claudin-5 isoform expression over time after stroke using integrated high-resolution 9.4 T MRI, sensorimotor assessment, and quantitative cerebrovascular protein analysis; (2) validate these findings through RNA sequencing analysis of both rodent and human ischemic cerebral tissue; and (3) identify potential novel splice variants of *CLDN5* that may contribute to BBB regulation in stroke pathophysiology.

## Methods

### Data Availability and Study Design

The data supporting findings are available from the corresponding authors upon reasonable request. Detailed methods, study design, and statistical analysis used are available in Supplemental Materials. Although females were not included in this study, we recognize the importance of studying both sexes, and future investigations are aimed at including male and female stroke models to identify sex differences utilizing intense imaging, neurologic tests, and molecular assays. All procedures and animal experiments were performed in compliance with the European Community Council Directive (2010/63/EU) for Protection of Vertebrate Animals Used for Experimental and other Scientific Purposes guidelines. The ethical permit (Animal Inspectorate License No. 5.8.18-10593/2020) was approved by the Malmö-Lund Institutional Ethics Committee under the Swedish National Department of Agriculture. Results generated are reported in compliance with the Animal Research: Reporting of *In Vivo* Experiments (ARRIVE) guidelines.

### Thromboembolic stroke model and treatment

Thromboembolic stroke was performed in twelve-week-old male Wistar rats (Janvier, France) as previously described.^27,28^ Rats were sedated with 3% isoflurane mixed in N_2_O/O_2_ (70:30) and then maintained at 1.5-2% for the duration of the surgery. A craniotomy was performed, exposing the middle cerebral artery (MCA) bifurcation after removing the dura. A laser Doppler flow meter was placed on the skull within the right MCA region to monitor cerebral blood flow (CBF). Thrombin (12 UI Nordic Diagnostica AB, Sweden) was injected into the MCA lumen using a micropipette, which remained in place for 20 minutes for clot stabilization. Successful surgery was confirmed by a 70% decrease in CBF, remaining stable for 1h. Animals were randomized into the following treatment groups: JNJ0966 (10mg/kg; selective MMP-9 inhibitor), BI-0115 (10mg/kg; selective oxidized low-density lipoprotein receptor 1 inhibitor), or the combination JNJ0966 plus BI-0115. Drugs were administrated i.p. at 3.5h post occlusion at which rt-PA 3mg/kg (Alteplase, Boehringer Ingelheim AB, Germany) or vehicle (saline) was given intravenously in the tail vein at 4h post occlusion starting with a bolus dose of 10 % followed by a 40min infusion.

### Magnetic resonance imaging (MRI)

MRI was used to evaluate infarct lesion size, edema, hemorrhage, and CBF at 24h post stroke onset. Animals were anesthetized with O_2_ mixed with 3% isoflurane which then was lowered to 1.5-2% for the duration of the imaging procedure. Imaging was performed using a 9.4T preclinical MRI horizontal bore scanner (Biospec 94/20, Bruker, Germany). T2-weighted images were acquired using the RARE sequence: 25 axial slices, slice thickness 0.8mm, 256x146 matrix, in-plane resolution 137x137µm^2^, TR = 270ms, TE = 33ms, bandwidth 33kHz, TA = 2min 25s. T2*-maps were reconstructed from a multi gradient-echo sequence acquired with parameters as above except: TR = 1800ms, TE = 3.5ms to 58.5ms in steps of 5ms, bandwidth 69kHz, TA = 3min 18s. Brain perfusion was measured using unbalanced Pseudo-continuous arterial spin labelling (pCASL) in PV6 using the implementation presented by Hirschler et al.^29^ In brief, anatomical images were acquired with a 2D RARE sequence with TE 33ms, TR 2,5s, resolution 117x117µm^2^, FOV 30x30mm and slice thickness 0.8mm and with 23 slices and two averages. Two control and label and phase optimization prescans were acquired to determine the phase settings for label and control in the pCASL scans. Labelling was applied in the rat’s neck −2cm from the isocenter during 1.5s followed by a post-labelling delay of 300ms. Pulses and gradients for labelling were set as in the original publication. Readout was reported with a 2D EPI sequence with TE 13ms, TR 2s, resolution 312x313µm^2^, FOV 24x30mm and slice thickness 4mm. Inversion efficiency was obtained as in the original publication with the exception that a resolution of 234 x 234µm^2^ was used with 4 averages. The transmission coil was used in transmit-receive mode as reconstruction of arrayed coil data was not possible for the inversion efficiency measurement in the current implementation of the method. For the pCASL perfusion measurement the original implementation was followed with the modification of TE = 14.2ms, TR = 4s, resolution 234x234µm^2^, FOV 18x29mm and slice thickness 2mm with 5 slices. Slices were placed to cover the stroke area otherwise centred around the isocentre. For the final T_1_ the modifications were TE=12.8ms and the geometry was taken from the preceding pCASL scan.

### Neurological evaluation

A 28-point neuroscore test and cylinder test prior to and 24h post stroke onset was used to assess neurological function as previously described.^28,30,31^ The 28-point neurological composite test was based on 11 different sensorimotor tasks.^31^ The animal’s performance of the various tasks was scored by an experienced evaluator blinded to the treatment groups. A score of 28-points, the accumulative score of the test, indicates a normal neurological function in a healthy rat.

### Tissue collection

Two different cohorts of animals were used for temporal profile studies and for drug treatment. Animals used for the temporal profile studies were euthanized 3, 6, and 24h post stroke and for drug treatment assessment, rats were euthanized 24h post stroke following MRI and neurological function assessment. During tissue collection, rats were heavily sedated with isoflurane before intracardiac perfusion with saline was performed. Brains were collected and snap-frozen before being stored at −80°C until further use for western blot. The cerebral vasculature and parenchyma from the whole brain isolated from sham and stroke induced animals were separated according to a previously described method.^28,32^

### Western Blot

Proteins of interest were evaluated in vessels and parenchyma fractions using our western blot protocol as previously described.^28,33^

### RNA-sequencing analysis of CLDN5 alternative splice isoform expression

To characterize the alternative splicing landscape of *CLDN5* in the context of ischemic stroke, we analyzed publicly available bulk RNA-sequencing datasets from human brain microvascular endothelial cells exposed to hypoxia (GSE163827) and a rat transient middle cerebral artery occlusion (tMCAO) model (GSE279377). These datasets provided transcriptomic profiles from ischemic brain regions in both species with an emphasis on the brain microvascular endothelium. Raw .sra files were downloaded locally using the NCBI SRA Toolkit (v.3.2.1) and subsequently converted to FASTQ format using the fasterq-dump utility. Initial quality control was performed using FastQC (v.0.11.9), followed by adapter trimming and base-quality filtering via Trim Galore (v.0.6.10). Trimmed reads were aligned to the species-specific reference genomes using STAR aligner (v.2.7.11b) in two-pass mode, enabling accurate detection of annotated and novel splice junctions. Genome assemblies and annotation files were obtained from Ensembl: GRCh38.p13 for human and mRatBN7.2 for rat. Aligned reads were processed with StringTie (v.2.2.1) for transcriptome assembly and isoform-level quantification. Transcript structures reconstructed by StringTie were used to cross-validate known and novel *CLDN5* isoforms and quantify transcript-level expression values in transcripts per million (TPM). For downstream comparison between rat and human *CLDN5* isoforms, genomic coordinates and exon-intron structures were aligned based on conserved synteny and Ensembl BioMart annotations. To detect and quantify splicing events, we used the MAJIQ (Modeling Alternative Junction Inclusion Quantification) pipeline (v.2.4). BAM files were processed with MAJIQ’s build module using curated GFF3 annotations ensuring correct transcript-parent relationships. Local splicing variations (LSVs) within the *CLDN5* gene locus were identified and quantified using MAJIQ’s psi module. LSV outputs were cross-referenced with StringTie transcript assemblies and TPM estimates. Splicing events and isoform expression profiles were visualized using Integrative Genomics Viewer (IGV) Sashimi plots and the IsoformSwitchAnalyzeR package. Functional domain annotations were overlaid using Ensembl BioMart to evaluate the potential structural and regulatory consequences of observed splice isoforms in the context of cerebrovascular injury.

### Statistical Analysis

The number of individual animals are referred to as “*n*” and P < 0.05 was considered statistically significant. Statistical analyses were performed utilizing either GraphPad Prism 10.5.0 or R 4.4.1 and data are represented with mean ± standard deviation. Data were assessed for normality utilizing Shapiro-Wilk tests and in instances where normal distribution were not met, non-parametric tests were employed as indicated in each figure legends. Multiple comparisons were performed with one-way or two-way ANOVA test with Tukey’s multiple comparisons post-hoc test and or direct comparisons with unpaired t-test. Precise p-values are reported. In addition, statistical comparisons between groups were also taken into consideration when performing the analyses. Correlative analyses were performed with simple linear regression. We acknowledge that any significant departure from normality not detected in this study using Shapiro-Wilk testing may be due to under power; however, ROUT analysis was performed at 1% to detect potential outliers that may influence parametric testing.

## Results

### Genome-wide Alternative Splicing Analysis Reveals CLDN5-Specific Splicing Events in the Rat tMCAO model

To characterize transcriptome-wide splicing alterations following transient middle cerebral artery occlusion (tMCAO), we performed RNA-sequencing analysis of cerebrovascular tissue from control and tMCAO rat models. RNA-seq reads were aligned to the to the Rattus norvegicus reference genome using hisat2 (v2.1.0) and splice junction events were quantified using Regtools. Hierarchical clustering analysis of heatmaps of log_2_-transformed junction counts for genes associated with four key stroke-relevant Gene Ontology categories: barrier function, inflammation, cell death, and oxidative stress reveal distinct splicing signatures between tMCAO and control conditions (**Fig 1A**). Notably genes involved in barrier function and inflammatory responses exhibited the most pronounced splicing alterations post stroke, suggesting coordinated post-transcriptional regulation of these pathways during cerebral ischemia. (**Fig 1A**). Given the critical role of claudin-5 in BBB integrity, we performed targeted analysis on the *CLDN5* locus. Samples were realigned to chromosome 11 using STAR and splice junctions were re-quantified with Regtools. A heatmap of log_2_ (event counts + 1) on chr11 shows a striking alteration in the alternative splicing events following tMCAO within the chromosomal loci (**Fig. 1B**). Comprehensive mapping of all detected splice junctions within the *CLDN5* genomic interval identified a previously unannotated splice event spanning chr11:82,213,199– 82,219,334 that is present in both the control and tMCAO samples (**Fig. 1C**). This novel junction contrasts the current annotation of *CLDN5*, suggesting that claudin-5 may undergo potential native alternative splicing which is conserved following tMCAO. To quantify transcript level expression changes, we applied IsoformSwitchAnalyzeR for differential transcript usage analysis. Although total *CLDN5* gene and isoform expression were reduced in tMCAO (left panels), the relative usage fraction of the predominant transcript ENSRNOT00000072027 remained unchanged (right), suggesting that the novel junction does not replace the predominant existing isoform (**Fig. 1D**).

**Figure 1.**
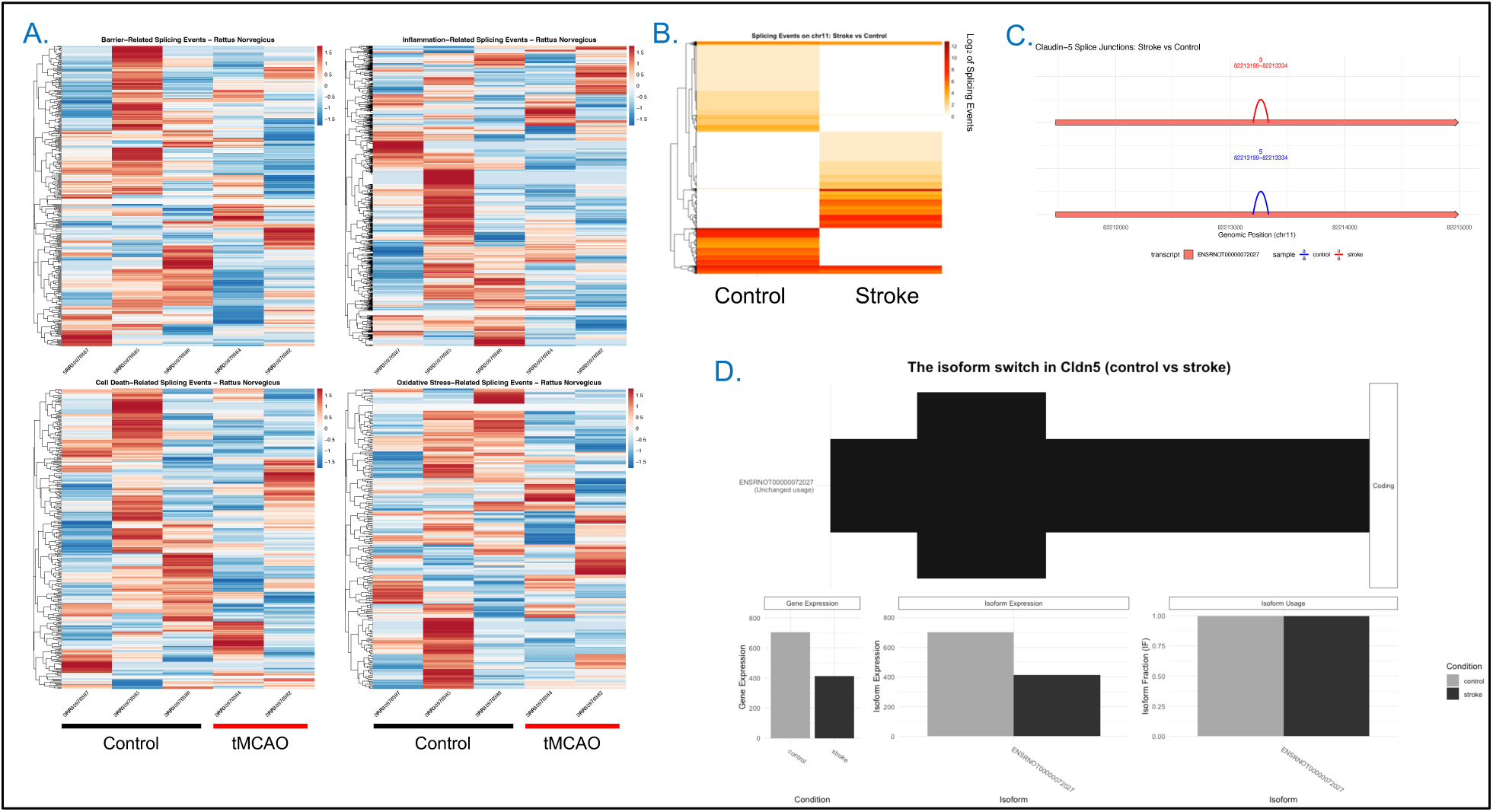
Genomic splicing profile of Rattus Norvegicus following tMCAO. (**A**) (**A**) Hierarchically clustered heatmaps of log_2_-transformed splice junction counts for genes in four stroke-relevant Gene Ontology categories: barrier function (top left), inflammation (top right), cell death (bottom left), and oxidative stress (bottom right). Each column represents an individual sample from either control or transient middle cerebral artery occlusion (tMCAO) rats. Distinct clustering patterns separate tMCAO from control samples, with the most pronounced splicing alterations observed in barrier function and inflammatory pathway genes, indicating coordinated post-transcriptional regulation of these pathways following ischemic injury. (**B**) Heatmap of log_2_(event counts) for all detected splicing events on rat chromosome 11 in control versus tMCAO samples. The marked reorganization of splicing patterns following stroke includes several loci showing coordinated differential junction usage, highlighting chromosome 11 as a hotspot for stroke-associated splicing alterations. (**C**) Comprehensive mapping of all detected splice junctions within the CLDN5 interval (chr11:82,213,199–82,219,334) reveals a previously unannotated splice event present in both control and tMCAO samples. This novel junction is absent from current annotations, suggesting that CLDN5 undergoes native alternative splicing that is conserved following cerebral ischemia. (**D**) IsoformSwitchAnalyzeR analysis of CLDN5 shows reduced total gene and isoform expression in tMCAO samples relative to controls (left and middle panels), while the relative usage fraction of the predominant transcript ENSRNOT00000072027 remains unchanged (right panel), indicating that the novel junction does not replace the primary annotated isoform.

### Validation of the Novel CLDN5 Splice Junction

To exclude potential technical artifacts and confirm that this novel CLDN5 junction truly reflects biological splicing we performed five orthogonal validation steps including: (1) splice motif validation, (2) comparison of RegTools vs. STAR, (3) transcript context, (4) SAMtools inspection, and (5) strand specificity. Splice site analysis revealed a non-canonical GC donor site paired with a canonical AG acceptor, consistent with legitimate, although rare, splice signals. While STAR’s default filters suppressed this junction, likely due to its low read support (3–5 reads), non-canonical donor, and minimal overhang, the junction was reproducibly detected by RegTools. The identified junction falls within the single canonical exon of *CLDN5*, indicating an intra-exonic splicing event not captured in current annotations. Manual inspection of BAM alignments revealed clean, uniquely mapped spliced reads spanning chr11:82,213,199–82,219,334, with no evidence of misalignment or sequencing errors. Strand specificity analysis of FLAG-tagged reads (SAM FLAGs 83/147/163/99) demonstrated overwhelming support for the negative strand, matching transcriptional orientation of *CLDN5* and ruling out antisense artifacts. Taken together, these validation steps confirm that the newly identified *CLDN5* splice event is a genuine, reproducible alternative junction, one that is absent from current Rattus norvegicus gene models and likely represents a low abundance but present transcript variant with potential functional significance.

### Temporal Dynamics of Claudin-5 Protein Isoform Expression Following Thromboembolic Stroke

To investigate the functional consequences of CLDN5 alternative splicing, we examined claudin-5 protein isoform expression in the isolated cerebrovasculature using our clinically relevant thromboembolic stroke model. Protein expression was analyzed at 3, 6, and 24h post-stroke onset in both ipsilateral and contralateral hemispheres (**Figs. 2A-C**). Via standard western blotting we observed multiple bands corresponding to the projected molecular weights of isoform 1 and isoform 2 claudin-5 (∼35kDa and ∼25kDa respectively) as well as a distinct lower band at ∼10kDa (Fragment) (**Fig. 2A**). Due to the relatively low expression of this ∼10kDa band we hypothesize that it may be a potential fragment of the identified isoform 2 which still expresses the epitope region of binding that the antibody is capable of binding to. Further investigation into the exact molecular composition of this identified structure is warranted but falls beyond the scope of this study. Following quantification of the detected claudin-5 isoforms there was an observed increase in isoform 1 of claudin-5 specifically within the ipsilateral hemisphere at 6h post-stroke onset (P=1.3×10^-2^) that was decreased at 24h (**Fig. 2B**). This observation of increased isoform 1 expression was not observed within the contralateral hemisphere (**Fig. 2C**). Interestingly, there were no corresponding changes to the expression of isoform 2 or fragment within both the ipsilateral and contralateral hemisphere (**Figs. 2B-C**). Indicating that an increase isoform 1 of claudin-5 could potentially be underlying alterations in cerebrovascular endothelial barrier integrity and in turn the permeability of the BBB following thromboembolic stroke.

**Figure 2.**
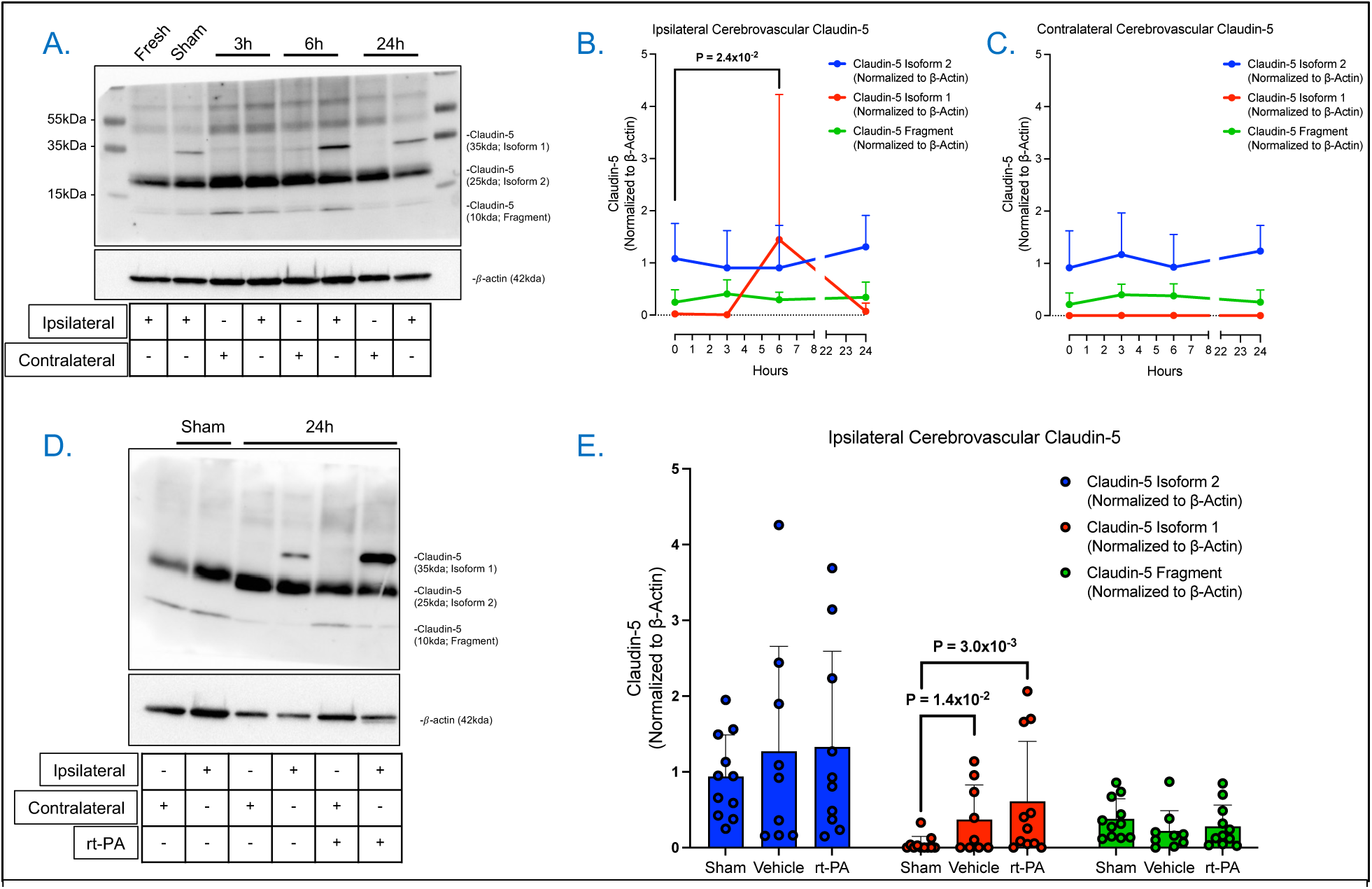
Temporal and rt-PA Mediated Claudin-5 Protein Isoform Expression Following Thromboembolic Stroke. (**A**) Representative image of western blot showing ipsilateral and contralateral cerebrovascular claudin-5 protein isoform expression following 3, 6, and 24h post-stroke onset as well as corresponding *β*-actin. (**B-C**) Densiometric quantification of claudin-5 isoforms normalized to corresponding *β*-actin at 3, 6, and 24h post-stroke in the isolated cerebrovasculature in the (**B**) ipsilateral and (**C**) contralateral hemispheres represented as the relative densities of claudin-5/*β*-actin. N=5-10 individual animals. Non-parametric Mann-Whitney test was employed to assess differences at 6h. (**D**) Representative image of western blot showing ipsilateral and contralateral cerebrovascular claudin-5 protein isoform expression following 24h post-stroke onset with or without delayed rt-PA treatment as well as corresponding *β*-actin. (**E**) Densiometric quantification of claudin-5 isoforms normalized to corresponding *β*-actin in sham and at 24h post-stroke treated with either vehicle or delayed rt-PA in the isolated cerebrovasculature in the ipsilateral hemisphere represented as the relative densities of claudin-5/*β*-actin. N=9-11 individual animals. Non-parametric Kruskal-Wallis test was employed to assess differences between treatment groups.

### Impact of Delayed Recombinant Tissue Plasminogen Activator Treatment Following Thromboembolic Stroke

Given previous observations that delayed rt-PA administration exacerbates brain edema, hemorrhage, and worsened neurological outcomes at 24h post-stroke onset^28^ we assessed claudin-5 expression at 24h post-stroke onset in the presence or absence of delayed rt-PA administration. Western blot analysis revealed the same claudin-5 expression profile as observed in the time-course study (**Fig. 2D**). While isoform 2 and the fragment expression remained unchanged following delayed rt-PA treatment (**Fig. 2E**) a subset of vehicle (n=4) as well as rt-PA treated animals (n=6) demonstrated an increase in isoform 1 expression, resulting in a statistically significant rise in isoform 1 compared to sham (**Fig. 2E**). These data in combination with our observations of delayed rt-PA mediated trend decreases in isoform 1 expression within the contralateral cerebrovasculature (P=8.0×10^-2^) (**Supplemental Fig. 1**) suggested that isoform 1 may be correlated to the worsened stroke outcomes that we observed previously. Therefore, we further interrogated the specific correlation of the different claudin-5 isoform expression with post-stroke outcomes within individual animals.

### Increased Claudin-5 Isoform 1 Expression Correlates with Cerebral Edema Formation

To elucidate the relationship between claudin-5 isoform expression and stroke outcomes we analyzed correlations between protein isoform levels and key outcomes measures including infarct volume, cerebral edema, neurological function and hemorrhagic transformation. To reveal the global relationships independent of treatment effects, we pooled data from all experimental animals regardless of experimental grouping and corresponding treatment (**Supplemental Fig. 2**). Correlation analysis between claudin-5 isoform expression and infarct volume revealed no significant associations for any of the three detected protein variants (isoform 1, isoform 2 and fragment) which is consistent with our previous observations^28^ that infarct size does not predict barrier dysfunction severity (**Figs. 3A-C**). In contrast, analysis of edema formation at 24 hours post-stroke revealed differential correlations among claudin-5 isoforms (**Figs. 3D-F**). While claudin-5 isoform 2 (**Fig. 3D**) and fragment (**Fig. 3F**) showed no significant correlation with edema. There was a positive correlation between claudin-5 isoform 1 and edema formation following stroke at 24h (**Fig. 3E**). These findings suggest that elevated isoform 1 expression may reflect a maladaptive vascular remodeling process that compromises rather than restores BBB integrity following ischemic injury.

**Figure 3.**
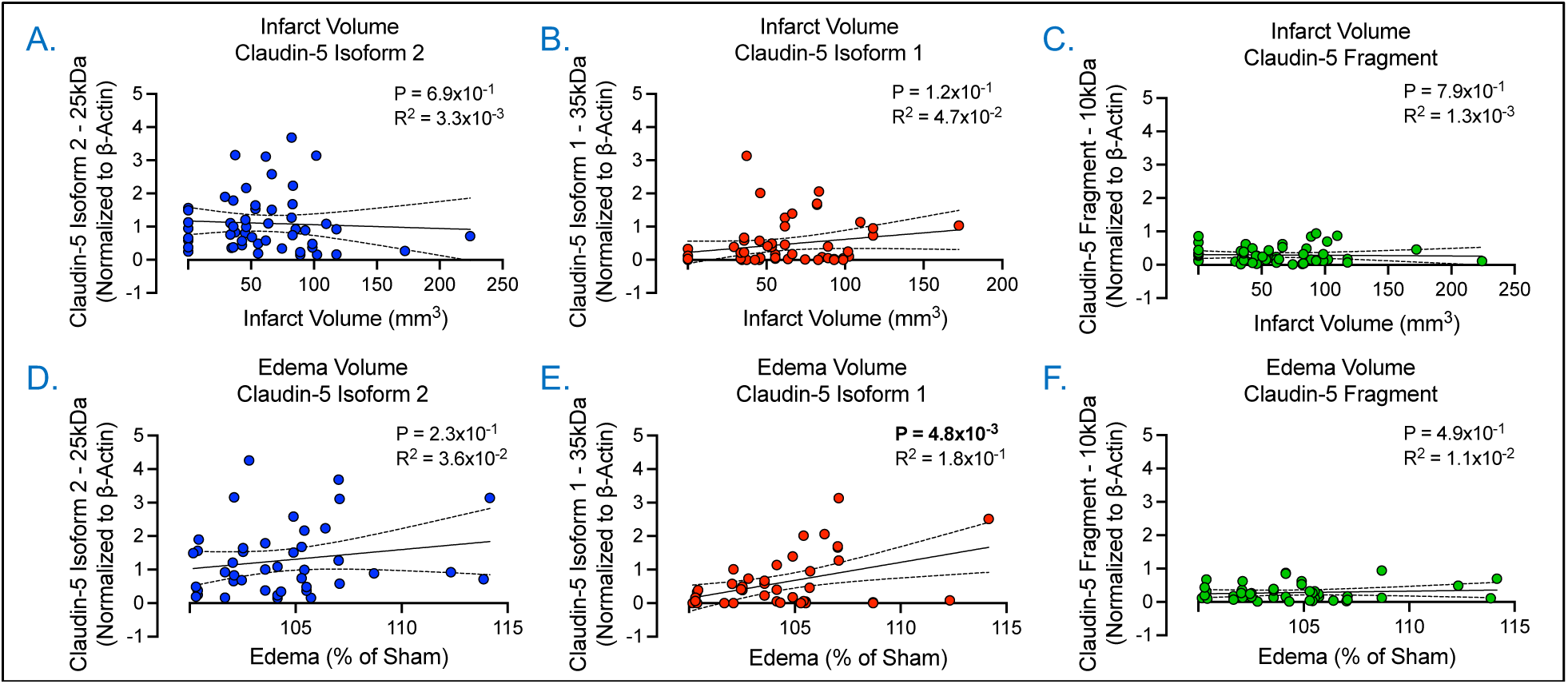
Correlation of Ipsilateral Claudin-5 Isoform Expression with Infarct Volume and Edema. (**A-C**) Graphical illustration of the individual correlation between (**A**) claudin-5 isoform 2, (**B**) claudin-5 isoform 1, and (**C**) claudin-5 fragment and infarct volume at 24h post-stroke onset. Simple linear regression was applied, and associated p-values as well as R squared values are shown within the graphical illustrations. N=52-54 individual animals. (**D-F**) Graphical illustration of the individual correlation between (**D**) claudin-5 isoform 2, (**E**) claudin-5 isoform 1, and (**F**) claudin-5 fragment and edema volume at 24h post-stroke onset. Simple linear regression was applied, and associated p-values as well as R squared values are shown within the graphical illustrations. N=41-44 individual animals.

### Claudin-5 Isoform 1 Correlates with Worsened Neurological Function and Hemorrhagic Transformation

Following our observations of claudin-5 protein isoform expression with infarct and edema volume (**Fig. 3**), we next sought to assess the relationship between these isoforms and neurological function (**Figs. 4A-C**) as well as hemorrhage (**Figs. 4D-F**). Correlation analysis revealed a significant positive association between increased expression of isoform 1 of claudin-5 (P=2.9×10^-3^) and neurological deficit severity at 24h post-stroke onset (**Fig. 4B**). This correlation with neurological function was not observed in either isoform 2 (**Fig. 4A**) or fragment claudin-5 (**Fig. 4C**) which supports our previous observations in this study. Counter to our hypothesis and our prior results, we observed a positive correlation between increased expression of both claudin-5 isoform 2 (P=4.5×10^-3^) (**Fig. 4D**) and isoform 1 (P=4.6×10^-3^) (**Fig. 4E**) and hemorrhage. In concordance with our previous observations, we did not observe a correlation between fragment claudin-5 and hemorrhage (**Fig. 4F**). The correlative observations made between both isoform 2 and 1 of claudin-5 were driven by the animals that exhibited severe hemorrhage but point to a potential underlying mechanism in which aberrant increased claudin-5 isoform 1 expression and possibly a compensatory upregulation of non-functional isoform 2 expression could be driving hemorrhagic transformation in severe strokes (R^2^=1.6×10^-1^). This is exemplified by the same analysis performed in which the animals with hemorrhage greater than 5mm^3^ were removed and revealed no correlation with either isoform (**Supplemental Fig. 3**). Together these data suggest that isoform expression of claudin-5 following ischemic stroke plays a key role in stroke outcomes and merits further investigation.

**Figure 4.**
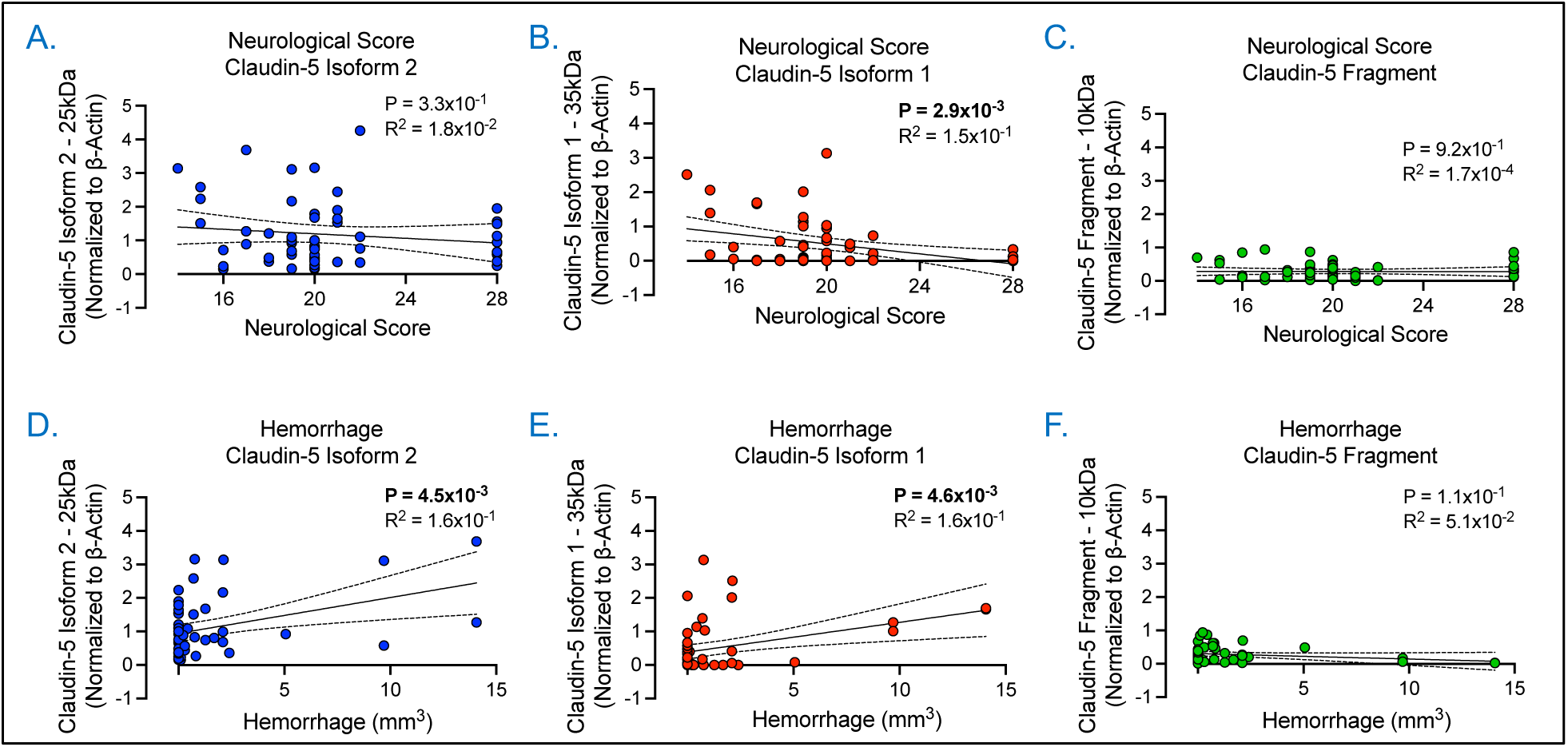
Correlation of Ipsilateral Claudin-5 Isoform Expression with Neurological Scores and Hemorrhage. (**A-C**) Graphical illustration of the individual correlation between (**A**) claudin-5 isoform 2, (**B**) claudin-5 isoform 1, and (**C**) claudin-5 fragment and 28-point neurological score at 24h post-stroke onset. Simple linear regression was applied, and associated p-values as well as R squared values are shown within the graphical illustrations. N=56-58 individual animals. (**D-F**) Graphical illustration of the individual correlation between (**D**) claudin-5 isoform 2, (**E**) claudin-5 isoform 1, and (**F**) claudin-5 fragment and hemorrhage volume at 24h post-stroke onset. Simple linear regression was applied, and associated p-values as well as R squared values are shown within the graphical illustrations. N=48-50 individual animals.

### Global and Claudin-5 Specific Splicing Alterations in Human Brain Endothelial Cells Following Ischemic-Like Injury

To investigate the impact of ischemic-like injury on alternative splicing, we performed chromosomal splicing analysis of cultured human brain endothelial cells exposed to hypoxia compared with controls. While no individual splicing events met chromosomal-wide statistical significance (**Fig. 5A**), the volcano plot and percent spliced in (PSI) distributions (**Fig. 5B**) reveal a clear shift in splicing patterns, suggesting that larger cohort sizes would reveal differential exon usage in response to injury. Clustering of chromosome 22 splicing events (**Fig. 5C**) further highlighted broad transcriptomic reorganization, including at the CLDN5 locus (**Fig. 5D**), a critical determinant of BBB integrity. Global splicing heatmaps identified condition-specific junction usage (**Fig. 5E**), and targeted junction analysis of CLDN5 (**Fig. 5F**, left) demonstrated differential transcript expression despite the absence of a statistically significant switch in isoform usage (P=0.567; **Fig. 5F**, right). Isoform structure analysis revealed four annotated CLDN5 transcripts (ENST00000403084.1, ENST00000406028.1, ENST00000413119.2, ENST00000618236.2), all exhibiting nearly identical expression profiles. TPM-based quantification showed high baseline expression in controls and a marked, uniform reduction across all isoforms in ischemic cells. This pattern indicates that ischemic-like injury triggers a broad transcriptional downregulation of CLDN5 rather than a selective shift in isoform abundance. The absence of a detectable isoform switch is consistent with the limitations of short-read RNA sequencing for resolving fine-grained splicing differences, particularly at the protein-coding C-terminal region where structural variation in isoform 1 is hypothesized to occur. We postulate that ischemic-like injury induces a shift toward increased isoform 1 protein expression of claudin-5, likely occurring at the post-transcriptional level, as supported by similar splicing alterations in both the human and rat genome.

**Figure 5.**
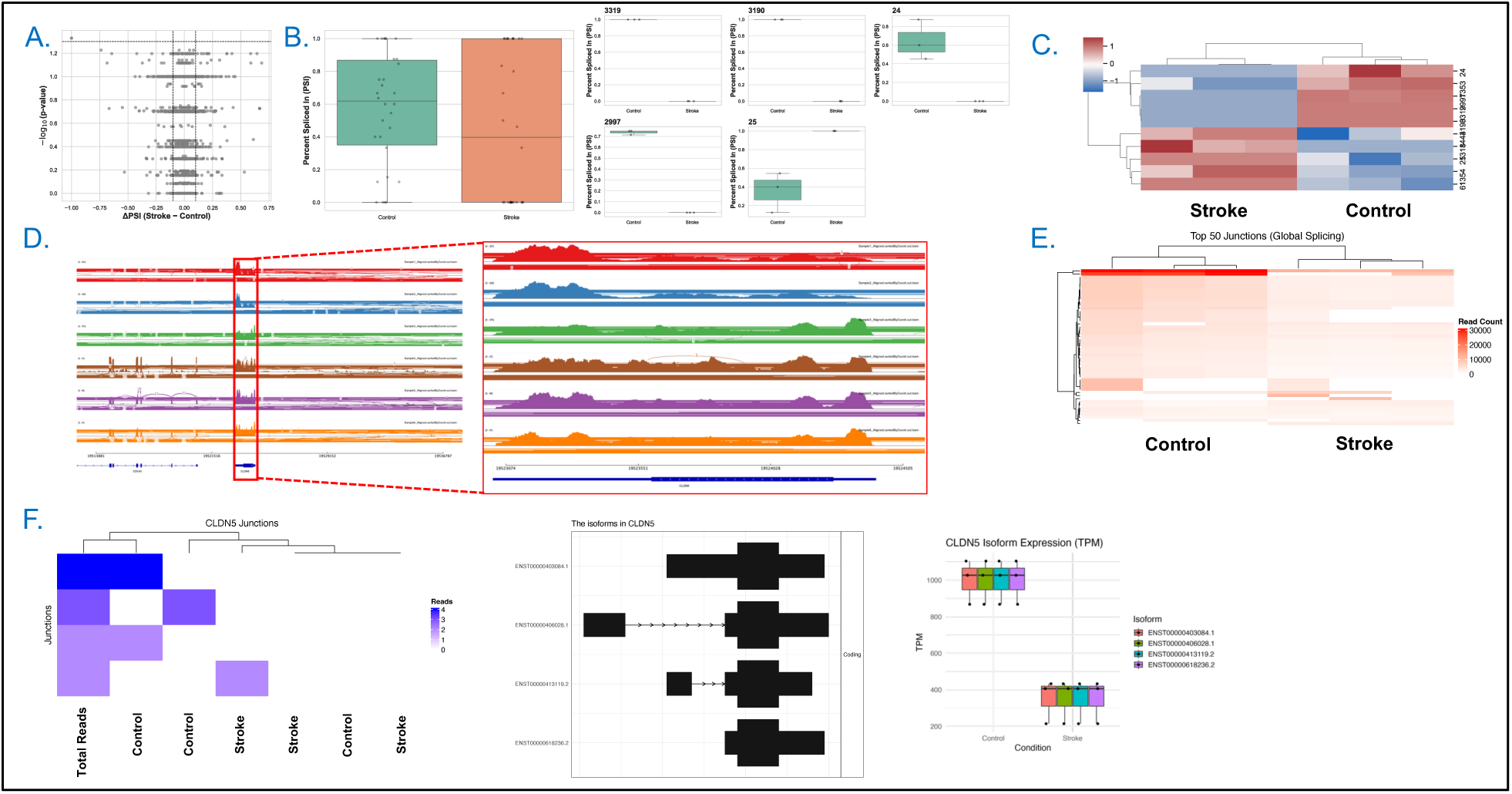
Integrative analysis of alternative splicing and junction usage reveals human brain endothelial splicing alterations following ischemic-like injury. (**A**) Volcano plot showing ΔPSI (Stroke − Control) on the x-axis versus −log_10_(p-value) on the y-axis for all detected splicing events on human chromosome 22. Red points denote significant events (adjusted p < 0.05, |ΔPSI| > 0.1), dashed lines indicate thresholds. (**B**) Distribution of Percent Spliced In (PSI) values for all events in control and stroke samples, with individual events overlaid. Right, PSI distributions for the top five identified differentially spliced events. (**C**) Hierarchically clustered heatmap of standardized PSI values for splicing events on human chromosome 22, showing distinct clustering of control versus stroke samples. (**D**) Integrative Genomics Viewer (IGV) sashimi plots of three control (top) and three stroke (bottom) samples. Left, expanded view of human chromosome 22 splicing patterns; right, magnified region encompassing CLDN5 highlighting altered junction usage and coverage in stroke. (**E**) Heatmap of the top 50 most highly supported splice junctions on chromosome 22 from STAR alignments, revealing global differences in junction usage between control and stroke groups. (**F**) Left, heatmap of CLDN5-specific junction read counts across all samples. Middle, predicted CLDN5 isoform transcript structures. Right, boxplots of CLDN5 isoform expression (TPM) showing markedly reduced expression in stroke relative to controls.

## Discussion

This study, characterize a previously unrecognized Claudin-5 splice isoform within the rodent cerebrovasculature as well as confirmed its presence in human brain microvascular endothelial cells following ischemic injury and demonstrate that its expression dynamics correlate with distinct stroke outcomes. Using a clinically relevant thromboembolic rat model along with analysis of both rodent and human RNA-seq datasets (GSE279377, GSE163827), we show that the 35kDa isoform 1 of CLDN5 is transiently upregulated in the ipsilateral cerebrovasculature at 6h post-stroke. Importantly, the observed elevated isoform 1 protein is positively associated with edema, worsened neurological scores, and hemorrhage, but not with infarct volume or cerebral blood flow (**Supplemental Fig. 4**), suggesting that this isoform primarily modulates BBB permeability and subsequently neurological outcomes rather than affecting neuronal survival directly. These observations are supported by recent research emphasizing the critical and potentially central role of cerebral edema on neurological outcomes following stroke.^34–36^ Additionally, secondary analysis of human and rat RNA-seq confirmed a conserved alternative splicing event within the CLDN5 locus, suggesting a post-transcriptional mechanism of BBB regulation in stroke accepted within the human annotated genome, that may also be present within the rodent cerebrovasculature.

The identification of alternative claudin-5 splicing provides novel insights into post-transcriptional mechanisms governing BBB integrity during cerebral ischemia. While disruption of endothelial tight junction integrity represents an established hallmark of ischemic injury, leading to edema and secondary neuronal damage^37–39^ our findings reveal a potentially critical mechanism underlying worsened stroke outcomes. Specifically, alternative splicing of claudin-5 in response to ischemia suggests that isoform identity within the tight junction complex can influence barrier function independently of cell-death pathways. Moreover, our findings suggest that detection of the previously identified 303 amino acid isoform within the human tissues is present within the rat cerebrovasculature and detectable via western blotting.

The identification of a distinct 35kDa isoform raises interesting questions about the structural and functional impact of splice-derived domain alterations. Previous studies have demonstrated that the c-terminus of claudins, including claudin-5, is critical for binding and stabilization of the zonula occludens complex^40–43^, which in turn regulates its functionality. We postulate that the extension of the c-terminus as observed in the 35kDa claudin-5 isoform in this study, may confer a similar action as compared to the phosphorylation of T207 via Rho-associated kinases which results in attenuated junctional localization and consequently function of claudin-5.^44^ Additional investigation into the sub-cellular localization is warranted; however, falls beyond the scope of this study. Our proposition of altered functionality of claudin-5 is supported by separate work which has shown that alternative inclusion or exclusion of extracellular loop sequences has the potential to modify paracellular pore size or ion selectivity.^45^ Moreover, rare de novo mutations within the CLDN5 coding sequence have been linked to neurological disease states, underscoring that even subtle alterations in CLDN5 structure can have profound effects on patient outcomes.^19,46^ We therefore posit the plausibility that isoform 1 represents a compensatory, and potentially maladaptive, form of CLDN5 that transiently integrates into the affected microvasculature which fails to restore full barrier competence. These findings suggest that claudin-5 isoform profiling may serve as a valuable biomarker for predicting stroke severity and treatment response.

Supporting the potential of claudin-5 as a stroke biomarker, several humans’ studies have established a link between circulating levels of claudin-5 protein and clinically relevant outcomes in stroke patients. In a prospective cohort study it was found that plasma levels of claudin-5 measured at 12h post-stroke independently predicted hemorrhagic transformation.^47^ These findings align with our current results in this study, further promoting the notion that deeper interrogation of claudin-5 at the level of isoform expression could yield more profound clinical insights. In contrast, a separate prospective cohort study found that circulating levels of both claudin-5 and occludin, an additional tight junction protein, did not exhibit a relationship to neurological status on the first day of stroke, but rather the anatomical location of the stroke.^48^ These results complement our findings, as we did not observe any significant correlation between isoform 2 of claudin-5, the predominantly expression protein isoform, and neurological function at 24h. It was only when stratified by isoform type that a clear relationship between claudin-5 and neurological function was established, thus further promoting the potential viability of utilizing claudin-5 isoform expression as a biomarker which may reveal underlying cerebrovascular injury dynamics. Further evidence comes from a study which included a broader panel of biomarkers, where patients that exhibited clinical deterioration due to hemorrhagic transformation had higher serum levels of claudin-5 within 3 hours of stroke onset.^49^ These findings in combination with our observations highlight the critical importance of timing in the evaluation of claudin-5 as a biomarker for stroke outcomes. Additionally, recent evidence indicates that serum claudin-5 levels can differentiate ischemic stroke from stroke mimics^50^, suggesting that isoform-specific assessment may provide enhanced diagnostic accuracy and sensitivity.

Beyond acute cerebrovascular stroke, claudin-5 isoform profiling may have broad utility as a biomarker across neurological disorders characterized by BBB dysfunction to determine disease progression and cognitive function. Potentially, expanding upon prior investigations which examined the relationship between plasma levels of claudin-5 within patients on the first day of acute ischemic stroke^48^, and including disease states such as multiple sclerosis, Alzheimer’s disease, and vascular dementia.^51^ Differential detection of the 35kDa versus 25kDa CLDN5 isoforms in patient cerebrospinal fluid or plasma-derived exosomes could provide enhanced specificity for identifying BBB breakdown.^52^ Furthermore, the involvement of CLDN5 splicing in chronic neurodegenerative and neuropsychiatric conditions, such as Parkinson’s disease and schizophrenia, is an exciting prospect that warrants systematic investigation.^53,54^ Understanding splicing regulation in these contexts may reveal novel therapeutic targets for maintaining blood-brain barrier integrity across diverse pathological conditions.

Advances in RNA-based therapeutics such as splice-switching oligonucleotides, have enabled this treatment modality to become a powerful and exciting avenue for potentially treating severe pathologies.^55,56^ These technologies enable precise regulation of pre-mRNA splicing mechanisms such as enhancers and silencers which have reached clinical development for diseases such as spinal muscular atrophy and Duchenne muscular dystrophy.^55^ Though direct application to ischemic stroke treatment remains promising^57–59^, studies emphasizing splicing dysregulation as a factor in stroke pathophysiology have revealed a potentially novel therapeutic angle.^60^ Given that our data shown here in this study identifies novel differential isoform 1 of claudin-5 expression, potentially secondary to unique splice switching, it is plausible to design oligonucleotides or other splice-modulatory agents to suppress the aberrant overexpression of this isoform which correlates with increased edema, hemorrhage, and poor neurological outcomes. Alternatively, this technology could be aimed at redirecting splicing towards the canonical isoform 2, to preserve tight junction integrity and mitigate BBB dysfunction. While challenges with regards to cell-type specificity off-target effects, and timing still pose a barrier, the field of RNA therapeutics is rapidly evolving and improving that could support a translational strategy to target maladaptive splicing mechanisms underlying claudin-5 isoform expression and potentially restore homeostatic generation of the key tight junction protein.

While our findings demonstrate significant correlations between claudin-5 isoform 1 expression and stroke outcomes, the mechanistic basis for these associations requires further investigation. Subcellular localization studies are needed to determine whether isoform 1 properly integrates into tight junction complexes or exhibits aberrant trafficking patterns. Additionally, functional studies examining the barrier properties of endothelial monolayers expressing different claudin-5 isoform ratios would provide direct evidence for our proposed mechanism. Future investigations should focus on identifying upstream splicing regulators that control claudin-5 isoform expression and determining their susceptibility to pharmacological modulation. Understanding the temporal dynamics of splicing regulation may reveal therapeutic windows for intervention to promote beneficial isoform expression patterns.

In summary, our work reveals that differential protein isoform expression potentially mediated via alternative splicing of Claudin-5 as a novel, conserved mechanism underlying BBB regulation in ischemic stroke. The preferential association of the novel isoform 1 with vasogenic edema and hemorrhage, but not infarct volume, highlights the importance of splicing-dependent fine-tuning of tight junction integrity and ultimately neurological function. These findings establish a foundation for developing claudin-5 isoform ratios as biomarkers for cerebrovascular disease and identify alternative splicing machinery as a potential therapeutic target for preserving BBB function in neurological disorders.

## Acknowledgments

The authors are grateful to Michael Gottschalk at Lund University Bioimaging Center (LBIC), Lund University for their assistance and providence of the magnetic resonance imaging facility.

